# A novel VCP modulator, KUS121, attenuates atherosclerosis progression by maintaining intracellular ATP and mitigating ER stress in endothelial cells

**DOI:** 10.1101/2024.04.29.591786

**Authors:** Fuquan Zou, Osamu Baba, Takahiro Horie, Yasuhiro Nakashima, Shuhei Tsuji, Tomohiro Yamasaki, Chiharu Otani, Sijia Xu, Miyako Imanaka, Kazuki Matsushita, Keita Suzuki, Eitaro Kume, Hidenori Kojima, Qiuxian Qian, Kayo Kimura, Naoya Sowa, Akira Kakizuka, Koh Ono

## Abstract

**Background:** Endoplasmic reticulum (ER) stress signaling pathways have pivotal roles in atherosclerosis progression. Recently, we have developed Kyoto University Substance (KUS) 121, which selectively inhibits ATPase activities of valosin-containing protein (VCP) and consequently saves intracellular ATP consumption and mitigates ER stress.

**Methods and Results:** We assessed the efficacy of KUS121 against atherosclerosis by its daily injection into *Apoe^−/−^* mice fed with Western Diet (WD) for 8 weeks. Consequently, KUS121 treatment reduced atherosclerosis progression by approximately 40% in atherosclerotic plaques. Interestingly, we found that C/EBP homologous protein (CHOP), an established ER stress marker, was mainly expressed in plaque endothelium. Therefore, we assessed the action of KUS121 in endothelial cells using the human endothelial cell line (EA.hy926 cells). As a result, KUS121 prevented ER stress-induced apoptosis and downregulated the IRE1 (Inositol-requiring enzyme) α-associated inflammatory pathways. Consistent with these in vitro findings, KUS121 treatment also significantly reduced endothelial apoptosis assessed by TUNEL staining and inflammation examined by immunostaining of Nuclear factor kappa B (NF-κB) and Intercellular adhesion molecule (ICAM) 1 at plaque endothelium. We also demonstrated that KUS121 maintained ATP levels in EA.hy926 cells and atherosclerotic plaque lesions using the single-wavelength or the FRET-based fluorescent ATP sensor. Supplementation of intracellular ATP by Methyl pyruvate (MePyr) attenuated ER stress-induced apoptotic and inflammatory pathways in endothelial cells, which could be the main mechanism how KUS121 reduces ER stress.

**Conclusions:** KUS121 can be a new therapeutic option for atherosclerotic diseases by maintaining intracellular ATP levels and attenuating ER stress-induced apoptosis and inflammation in plaque endothelium.

## Introduction

Atherosclerotic diseases such as cerebral infarction and coronary artery disease caused by stenosis or occlusion of arteries are the principal cause of mortality all over the world.^1,2^ Over the past few decades, the management of risk factors for atherosclerosis such as hypertension, dyslipidemia and diabetes has been advanced, and new treatment technologies for atherosclerosis such as stent implantation have been developed. However, there is still no treatment option for atherosclerosis, which directly affect systemically developed atherosclerotic plaques.

The pathophysiology of atherosclerosis is a complex process and affected by several factors.^3,4^ Among those, recent evidence has been increasingly suggesting that endoplasmic reticulum (ER) stress plays a crucial role in the development of atherosclerosis. Various atherosclerosis-inducing factors, including hyperlipidemia, oxidative stress, and low shear stress, disturb ER homeostasis, subsequently triggering ER stress.^4–7^ Chronic ER stress is thought to influence the formation of atherosclerotic plaques via multiple mechanisms. Apoptotic and inflammatory pathways are involved in the downstream of ER stress responses, and accelerate plaque formation and vulnerability ^5,8–10^.

Kyoto University Substance (KUS) 121 is a small molecule compound specifically designed to inhibit the ATPase activity of valosin-containing protein (VCP). VCP, which belongs to the AAA (ATPases Associated with diverse cellular Activities) family, is a ubiquitously expressed protein found across essentially all cell types. It assumes to perform a range of functions, including the cellular proteasome system, protein degradation related to the endoplasmic reticulum (ER), and facilitating membrane fusion.^11,12^ KUS121 does not disturb these cellular functions of VCP,^13^ but only inhibits its ATP consumption, which indicates that KUS121 works as a “VCP modulator”. In many stressed conditions, KUS121 have been demonstrated to prevent the decline in intracellular ATP levels by inhibiting the ATPase activity of VCP. It has been also demonstrated that KUS121 reduces ER stress and subsequent cell death, and these benefits are expected to be couple with the maintenance of intracellular ATP. ^13^

KUS121 showed its efficacy in preventing cell death across various animal disease models, which include several ophthalmic diseases, ischemic stroke and neurodegenerative disorders such as Parkinson’s disease.^13–19^ Furthermore, its Phase I/II clinical trials have indicated its safety of local administration and potential effectiveness in patients with non-arteritic central retinal artery occlusion.^20^ Our group also found that KUS121 reduced infarcted areas in ischemia and reperfusion injury models, and ameliorated cardiac functions in acute and chronic heart failure models.^21,22^

Here, we demonstrate that KUS121 is able to attenuates atherosclerosis progression in *Apoe^−/−^* mice. Unexpectedly, we found that ER stress primarily occurred at plaque endothelium. KUS121 effectively preserved intracellular ATP levels and suppressed ER stress responses in endothelial cells, which in turn mitigated endothelial apoptosis and inflammation both *in vitro* and *in vivo*. These results suggest not only that KUS121 is a promising candidate for the treatment of atherosclerosis but also that maintenance of intracellular ATP levels in endothelium can be a new therapeutic target of atherosclerosis via attenuation of ER stress.

## Methods

### Mice

*Apoe^−/−^* (B6.129P2-Apoetm1Unc/J, 002052) mice were purchased from Jackson Laboratories. Mice were fed with normal chow until 6 weeks of age. Then, diet was switched to Western diet (WD) (Research Diets Inc) containing 0.21% cholesterol and 41% milk fat to accelerate atherosclerotic plaque formation. To evaluate the impact of KUS121 on the progression of atherosclerotic plaques, mice were administered a daily intraperitoneal injection of 50 mg/kg KUS121, starting at 6 weeks of age. Control mice were injected a 5% glucose solution. Mice were sacrificed and samples were collected after 8 weeks of KUS121 treatment (Figure 1A). Mice were accommodated in specific pathogen-free animal facilities maintained by the Institute of Laboratory Animals within Kyoto University Graduate School of Medicine. All experiment procedures were approved by the Ethic Committee for Animal Experiments of Kyoto University.

**Figure 1.**
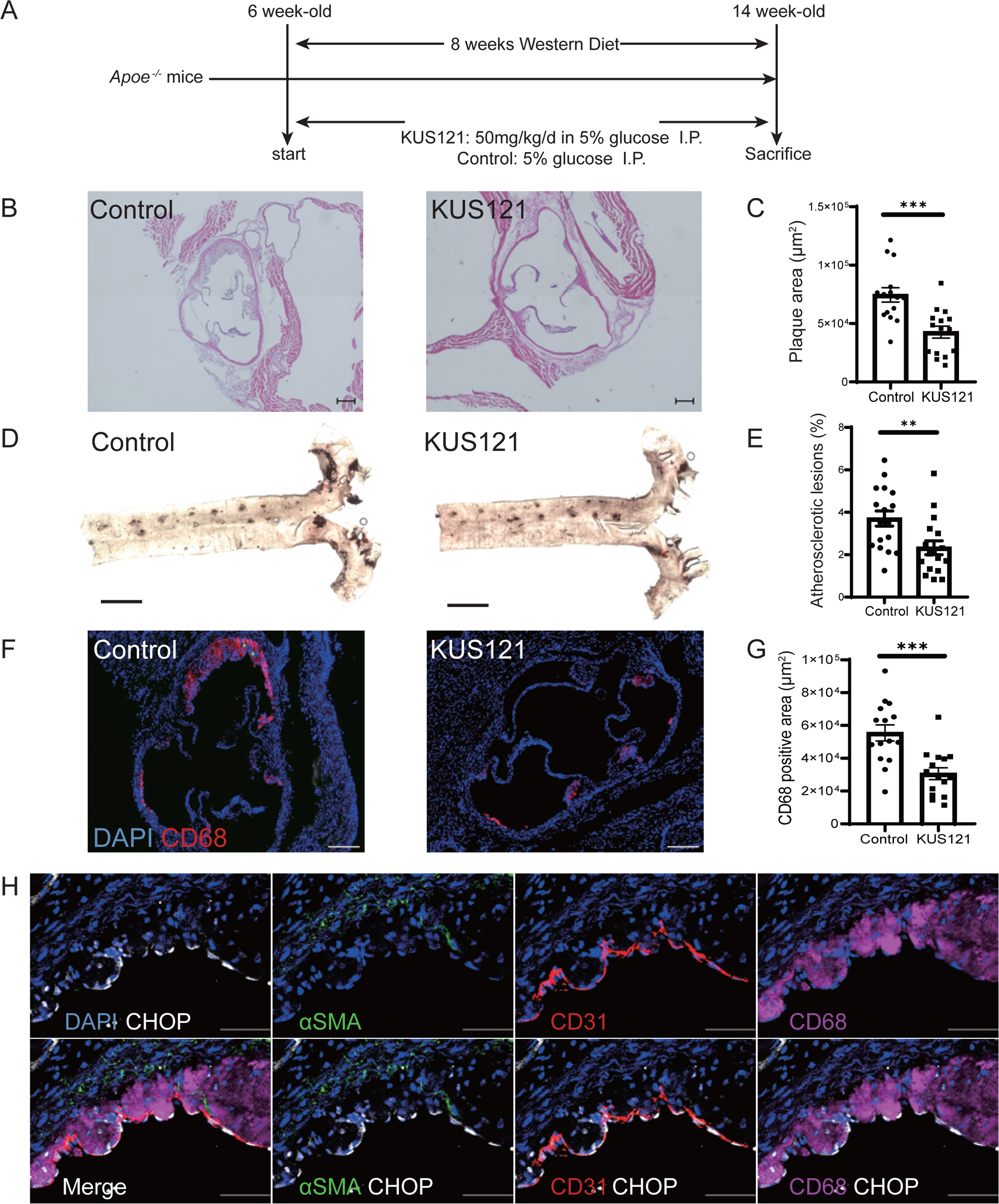
KUS121 attenuated atherosclerosis progression *in vivo*. **(A)** Scheme of experimental protocol to assess the effect of KUS121 on atherosclerosis progression. **(B)** Representative images of HE staining in aortic sinuses and **(C)** quantitative analysis of plaque areas (n=15 per group). Scale bar: 200 μm. **(D)** Representative images of en face ORO staining of whole thoracic aorta and **(E)** quantification of atherosclerotic lesions (n=17 per group). Scale bar: 2 mm. **(F)** Representative images of CD68 staining in aortic sinuses and **(G)** quantification of CD68-positive areas in atherosclerotic plaques (n=15 per group). Scale bar: 200 μm. **(H)** CHOP staining at the atherosclerotic plaque. Scale bar: 50 μm. All data are presented as mean ± SEM **p<0.01, ***p<0.001 by student’s *t* test.

### Quantification of Atherosclerotic plaque lesions

Atherosclerotic lesions were quantified by Hematoxylin and Eosin (HE) (Wako) staining of the cryosections of the aortic sinus and by en face analysis of the whole thoracic aorta stained by oil red O (ORO).^23–25^ After sacrifices, hearts were fixed and embedded in OCT compound for cryosections (10µm) of aortic sinuses. Sections were then subjected to HE staining for quantification. The average plaque size of 3 sections from each mouse was used as a representative value. For en face analysis, aortas were fixed and pretreated with 100% propylene glycol (Sigma-Aldrich) for 15 min, then incubated with ORO solution (Sigma-Aldrich) for 3 h at room temperature (RT). They were washed with 85% propylene glycol and then with PBS. After staining, aortas were placed between a cover glass and a slide grass, and imaged by LSM 710 (ZEISS). Images were analyzed using Image J software.

### Cell Culture

EA.hy926 cells (CRL-2922™, ATCC) and THP-1 derived macrophages were cultured in Dulbecco’s modified Eagle’s medium (DMEM) and RPMI 1640 (Nacalai Tesque) supplemented with 10% FBS and antibiotics (Gibco). To induce endoplasmic reticulum (ER) stress, 1 µg/mL tunicamycin (Sigma-Aldrich) was added to the cultured cells. Cells were also treated with KUS121, Methyl pyruvate (Wako) or Adenosine triphosphate (ATP) (Jena Bioscience) depending on the experiment.

### Assessment of cell viability in vitro

EA.hy926 cells were plated in a 96-well plate, followed by incubating with tunicamycin and KUS121 for 48 h. Then, cell viability was assessed by Cell Counting Kit-8 (Dojindo) according to the instruction manual. The absorbance value was measured by 2030 ARVO X (PerkinElmer).

### Quantitative Real-time PCR

Total RNA was isolated and purified using TriPure Isolation Reagent (Roche), and cDNA was synthesized from 1 µg of total RNA using Verso cDNA Synthesis Kit (Thermo Scientific) in accordance with the manufacturer’s instructions. For quantitative real-time PCR (qPCR), specific genes were amplified with THUNDERBIRD® SYBR qPCR Mix (TOYOBO) using StepOnePlusTM (Thermo Scientific). Expression levels were normalized using housekeeping gene, *ACTB*. The primer sequences are as follows:

*ACTB* forward, 5’-AGGCACTCTTCCAGCCTTCC-3’;
*ACTB* reverse, 5’-GCACTGTGTTGGCGTACAGG-3’;
*CHOP* forward, 5’-AGCTGTGCCACTTTCCTTTC-3’;
*CHOP* reverse, 5’-CAGAACCAGCAGAGGTCACA-3’;
*BIP* forward, 5’-TGTTCAACCAATTATCAGCAAACTC-3’;
*BIP* reverse, 5’-TTCTGCTGTATCCTCTTCACACGT-3’;
*sXBP* forward, 5’-CTGAGTCCCGAATCAGGTGCAG-3’;
*sXBP* reverse, 5’-ATCCATGGGGAGATGTTCTGG-3’;
*ATF4* forward, 5’-GTTCTCCAGCGACAAGGCTA-3’;
*ATF4* reverse, 5’-ATCCTGCTTGCTGTTGTTGG-3’;
*IL6* forward, 5’-CCTGAACCTTCCAAAGATGGC-3’;
*IL6* reverse, 5’-TTCACCAGGCAAGTCTCCTCA-3’;
*IL8* forward, 5’-ACTGAGAGTGATTGAGAGTGGAC-3’;
*IL8* reverse, 5’-AACCCTCTGCACCCAGTTTTC-3’;
*IL1B* forward, 5’-AGCTACGAATCTCCGACCAC-3’;
*IL1B* reverse, 5’-CGTTATCCCATGTGTCGAAGAA-3’;
*ICAM1* forward, 5’-GGCTGGAGCTGTTTGAGAAC-3’;
*ICAM1* reverse, 5’-ACTGTGGGGTTCAACCTCTG-3’;

### Western Blotting

Western blotting was performed using standard procedures as described previously.^26^ A total of 15 µg of protein was fractionated using NuPAGE 4–12% Bis-Tris Mini gels (Thermo Scientific) and transferred to Protran nitrocellulose transfer membranes (Whatman). The membrane was blocked using PBS containing 5% non-fat milk or 5% bovine serum albumin (BSA) for 30 min and incubated with the primary antibody overnight at 4°C. After washing by PBS containing 0.05% Tween-20 (PBST), the membrane was incubated with the secondary antibody for 1 h at RT. The membrane was washed again with PBST and then Pierce^TM^ Western Blotting Substrate (Thermo Scientific) or Pierce^TM^ Western Blotting Substrate Plus (Thermo Scientific) was used to detect proteins by LAS-4000 Mini system (Fuji Film). The following primary antibodies were purchased and used: anti-VCP (7F3, Cell signaling technology, #2649), anti-BIP (Cell signaling technology, #3183), anti-CHOP (L63F7, Cell signaling technology, #2895), anti-p-JNK (Thr183/Tyr185) (81E11, Cell signaling technology, #4668), anti-JNK (Cell signaling technology, #9252), anti-p-NF-κB (Ser536) (93H1, Cell signaling technology, #3033), anti-NF-κB (D14E12, Cell signaling technology, #8242), anti-p-IRE1α (Ser724) (Novus, NB100-2323), anti-IRE1α (14C10, Cell signaling technology, #3294), anti-ACTB (AC-74, Sigma-Aldrich, A5316). The secondary antibodies are as follows: anti-Rabbit IgG, HRP-Linked (GE Healthcare, NA934V), anti-Mouse IgG, HRP-Linked (GE Healthcare, 931V).

### Immunofluorescence Staining

For immunofluorescence staining, sections were blocked with 3% BSA in the presence of 1% Triton X-100 for 1 h at RT. Then, sections were incubated with primary antibody overnight at 4°C. After washing with PBS, sections were incubated with secondary antibody for 90 min at RT. Nuclei were stained using mounting medium with DAPI (Fluoro-Gel II Mounting Medium, Electron Microscopy Sciences). Immunofluorescence images were acquired using BZ-X810 all-in-one fluorescence microscope (Keyence), LSM 710 (ZEISS) or TCS SP8 microscope system (Leica). Images were analyzed using Image J and Imaris software (Oxford Instruments). Following antibodies were purchased and used for immunostaining: CD68 (FA-11, Bio-Rad), Actin, α-Smooth Muscle (1A4, Sigma-Aldrich), CD31 (2H8, Merck Millipore), CD31 (MEC13.3, BD Pharmingen), ICAM1 (YN1/1.7.4, BioLegend), CHOP (F-168, Santa Cruz), NF-κB (C-20, Santa Cruz). TUNEL staining was performed using In-Situ Cell Death Detection Kit (Roche Applied Science), according to the manufacturer’s instructions. The sections were co-stained with CD31 to identify Endothelial cells. The number of apoptotic endothelial cells (TUNEL^+^ and CD31^+^ cells) was counted and divided by the total number of endothelial cells.

### Serum data

Blood was obtained by submandibular vein puncture after 8 weeks of KUS121 treatment. Serum was separated by centrifugation at 4 °C and stored at −80 °C. Biochemical examination was performed using Hitachi 7180 Auto Analyzer (Nagahama Life Science Laboratory, Nagahama, Japan).

### Monocytes Recruitment

Monocyte recruitment was assessed by counting bead-labeled monocytes in atherosclerotic plaques as previously described^27^. Briefly, 200 µl of clodronate liposomes (Liposoma BV) were intravenously administered 5 days before sacrifice, and then, Ly6C^hi^ monocytes were labeled by intravenous injection of 1.0 µm Fluoresbrite® YG Microspheres (Polysciences, Inc) 2 days before sacrifice.

### Complete Blood Count and Flow Cytometry analysis

Peripheral blood was taken by submandibular vein puncture after 7 weeks of KUS121 treatment for the complete blood count and the flow cytometry analysis. Complete blood cell counts were measured using a hematology cell counter (Celltac α, NIHON KOHDEN). In preparation for flow cytometry, red blood cells were lysed using a commercial RBC lysis solution (BD PharmLyse, BD Biosciences). Samples were then incubated with antibodies for 30 min at 4°C. Following antibodies were purchased and used for flow cytometry analysis: CD45 (30F-11, BD bioscience and Biolegend), CD11b (M1/70, Biolegend and eBioscience), Ly6G (1A8, Biolegend), CD115 (AFS98, eBioscience), Ly6C (HK1.4, Biolegend), B220 (RA3-6B2, Biolegend), CD3ε (145-2C11, eBioscience).

For the *in vitro* apoptosis assay, EA.hy926 cells were stained with Alexa Fluor 647-Annexin V (BioLegend) and Helix NP^TM^ Green (BioLegend) at RT for 15 min in the dark for flow cytometry analysis. The samples were analyzed using a BD FACSAria II flow cytometer.

### Qualification of intracellular ATP levels in vitro

The single-wavelength fluorescent sensor (iATPSnFR^1.0^) was transduced into EA.hy926 cells using lentiviral transfection.^28^ ER stress was induced by 1 µg/mL tunicamycin (Sigma-Aldrich) and the cells were also incubated with KUS121 or 5 % glucose solution as control. After 48 h of incubation, the cells were washed with PBS and fluorescence images were taken by LSM 710 (ZEISS). The cells were also incubated with Methyl pyruvate (MePyr) or 2-Deoxy-D-glucose (2DG) depending on the experiments. The fluorescence intensity of the ATP sensor was analyzed using Image J. The intracellular ATP levels were also assessed by flow cytometry. In brief, to eliminate the influence of detachment of cells on intracellular ATP levels, the iATPSnFR^1.0^ transduced cells were cultured in floating state for 6 h with 1 μg/mL tunicamycin and KUS121 at 37 °C. Before flow cytometry analysis, the cells were washed with PBS and resuspended in PBS supplemented with 2 % FBS and 1 mM of EDTA. FlowJo (BD Biosciences) was used to analyze the fluorescence intensity.

### Quantification of ATP levels at atherosclerotic plaques using Go-ATeam2 mice

Go-ATeam2 mice expressing FRET-based ATP biosensor fluorescent protein reporters (orange fluorescent protein (OFP) and green fluorescent protein (GFP)) were kindly given by Dr. Masamichi Yamamoto.^22^ To assess ATP levels at atherosclerotic plaques, Go-ATeam2 mice were crossed with *Apoe^−/−^* mice and fed with WD for 12 months. The mice were anesthetized with intraperitoneal injection of 3 types of mixed anesthetic agents (medetomidine; 0.3 mg/kg, midazolam; 4.0 mg/kg, butorphanol; 1.0 mg/kg). Then, the common carotid artery was exposed, and fluorescence images of FRET and donor channels were taken by Olympus SZ16 microscope with the attached Reflected Fluorescence Illuminator (SZX2-RFA16). The OFP/GFP emission ratios were analyzed using Image J.

### Statistical Analysis

Measured data were presented as mean ± standard error of the mean (SEM). For statistical comparisons between 2 groups, Shapiro-Wilk test was performed to assess the normality of distribution. When the data was normally distributed, unpaired Student’s *t* test was conducted. Otherwise, Mann-Whitney *U* test was conducted. For statistical analysis of three or more groups, 1-way analysis of variance (ANOVA) with the Tukey’s multiple comparisons test was used. To analyze the time-dependent graph, 2-way ANOVA with the Tukey’s multiple comparisons test was performed. A p-value of <0.05 was considered as statistically significant.

## Results

### KUS121 Attenuates Atherosclerotic Progression in Apoe^−/−^ mice

At first, we assessed the effect of KUS121 on atherosclerosis progression using *Apoe^−/−^* mice fed with WD (Figure 1A). The daily administration of KUS121 significantly attenuated atherosclerosis formation at aortic roots (Figure 1B-C) and thoracic aortas (Figure 1D-E). Macrophage accumulation assessed by CD68 staining was also reduced by KUS121 treatment (Figure 1F-G). We also analyzed peripheral blood cell counts in those 2 groups. Interestingly, we observed a significant reduction in the number of peripheral neutrophils and Ly6C^hi^ inflammatory monocytes in the KUS121-treated group (Figure S1). On the other hand, there were no significant differences in body weight and chow consumption between the control and KUS121-treated groups (Figure S2A and B). Moreover, KUS121 treatment did not affect the serum profile including the cholesterol levels (Table S1).

As mentioned above, KUS121 has been demonstrated to reduce the ER stress.^13^ Then, we assessed the distribution of ER stress at atherosclerotic plaques in *Apoe^−/−^* mice on WD for 8 weeks by the staining of C/EBP homologous protein (CHOP), which is a well-known marker of ER stress, and found that CHOP was primarily expressed by plaque endothelial cells. Macrophages also expressed CHOP, but at much lower levels compared to endothelial cells. On the other hand, smooth muscle cells barely expressed CHOP (Figure 1H and S2C). These findings indicate that endothelial cells can be a potential primary target of KUS121.

### KUS121 protects endothelial cells from apoptosis under ER Stress

ER stress is known to trigger apoptosis, contributing to the progression of atherosclerosis.^7^ Thus, we evaluate the impact of KUS121 on cellular viability and apoptosis of endothelial cells. An ER stress inducer, tunicamycin reduced the viability of a human endothelial cell line, EA.hy926. However, this effect is reversed by KUS121 in a dose-dependent manner (Figure 2A-B). Then, we assessed the anti-apoptotic effect of KUS121 in endothelial cells by the Annexin V-binding assay. As expected, KUS121 treatment showed a dramatic reduction in early apoptotic cells (Annexin V^+^ Helix Green^−^) (Figure 2C-D). Moreover, the dead cells (Annexin V^+^ Helix Green^+^) were also significantly decreased (Figure 2E), which is consistent with the result of the viability assay in Figure 2A-B.

**Figure 2.**
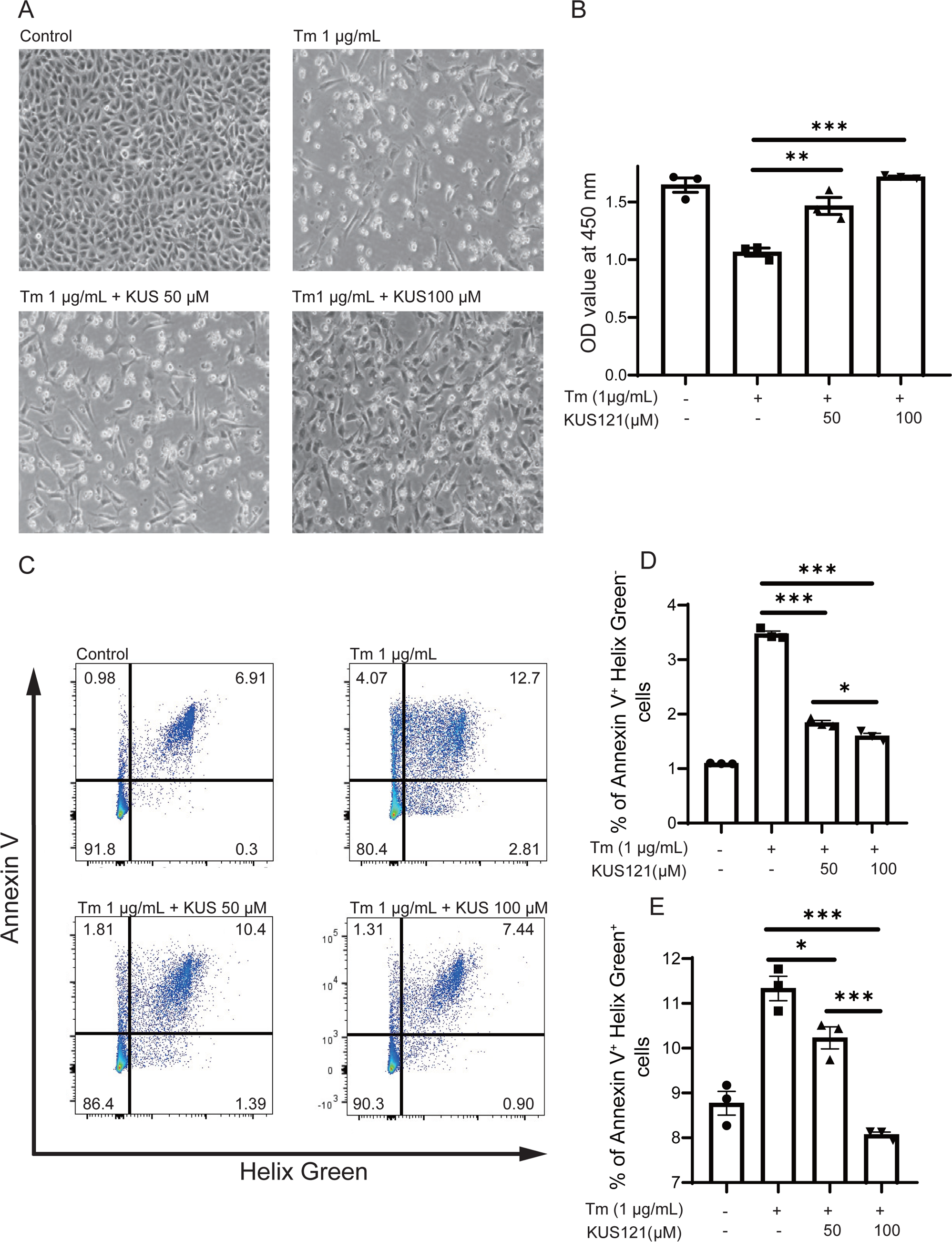
KUS121 protected endothelial cells from apoptosis. **(A)** Representative images of EA.hy926 human endothelial cells cultured with 1 μg/mL tunicamycin (Tm) for 48 h treated with different concentrations of KUS121 (50 and 100 μM) or DMSO as control and **(B)** quantification of cell viability (n=3 per group). **(C)** Representative dot plots of tunicamycin-induced apoptosis in EA.hy926 cells treated with different concentrations of KUS121 (50 and 100 μM) or DMSO as control. Quantification of **(D)** early apoptosis (Annexin V^+^ Helix Green^−^) and **(E)** cell death (Annexin V^+^ Helix Green^+^) (n=3 per group). The experiments were independently repeated 3 times. All data are presented as mean ± SEM. *p<0.05 **p<0.01, ***p<0.001 by 1-way ANOVA with the Tukey’s multiple comparisons test.

### KUS121 can attenuate both apoptotic and inflammatory pathways of ER Stress in endothelial cells

Next, given the central role of ER stress in plaque formation,^29,30^ we endeavored to further substantiate the effect of KUS121 on ER stress. To this end, we examined the effect of KUS121 on the signaling pathways of ER stress using EA.hy926 cells treated with tunicamycin. As anticipated, KUS121 effectively mitigated the upregulation of key genes in an ER stress response, such as Activating Transcription Factor 4 (ATF4), CHOP, Binding Immunoglobulin Protein (BIP), and spliced X-Box Binding Protein 1 (sXBP1) at mRNA levels (Figure 3A). To further validate the effect of KUS121 attenuating ER stress-mediated apoptosis, we assessed the protein expressions of CHOP and BIP which are the direct upstream of apoptosis-inducing signaling in the ER stress response and confirmed they were also reduced by KUS121 treatment. On the other hand, KUS121 treatment did not change the expression of VCP itself (Figure 3B-C).

**Figure 3.**
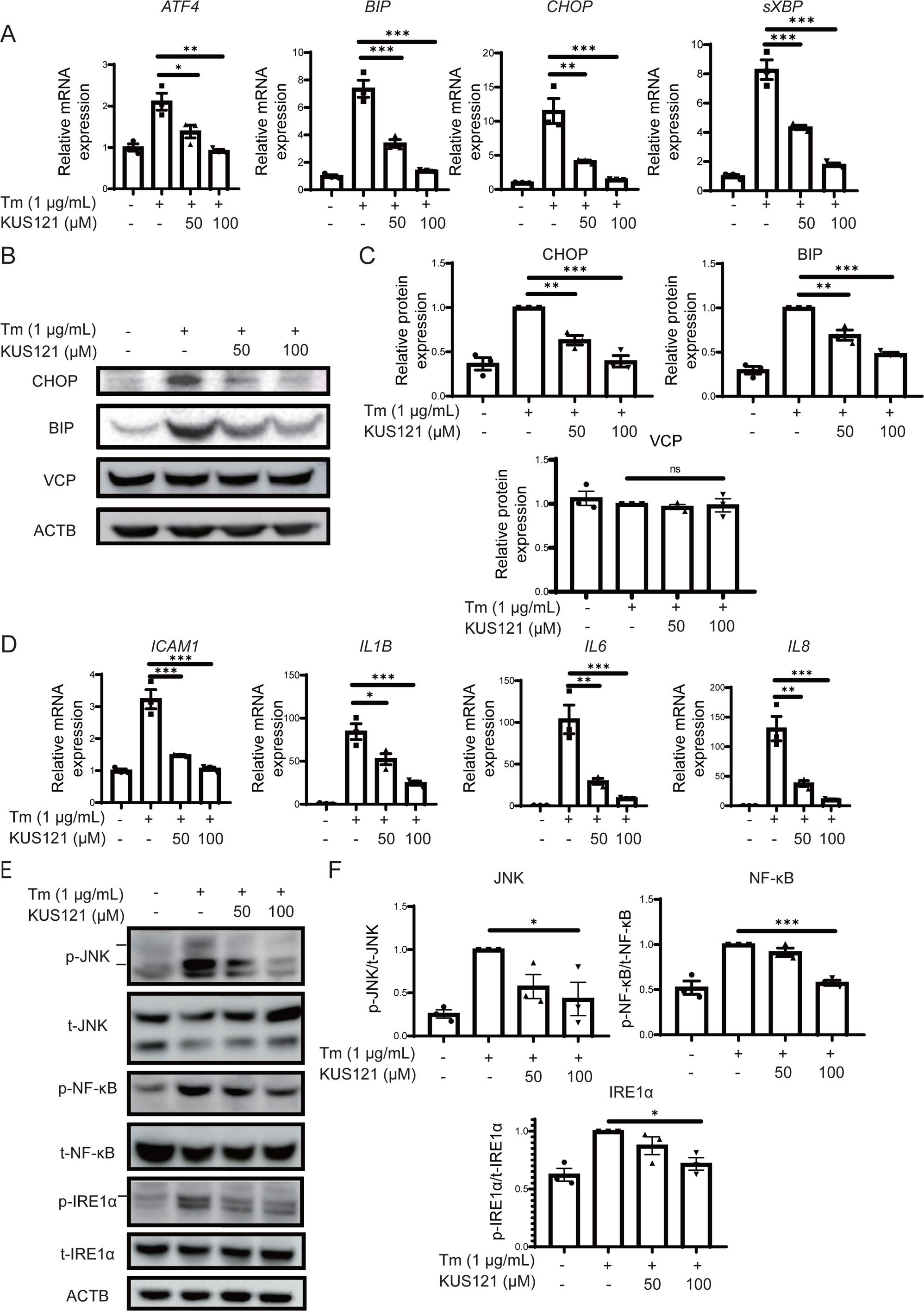
KUS121 mitigated not only the ER stress-induced apoptosis pathway but also the inflammatory pathways in human endothelial cells. **(A)** Quantitative real-time PCR analyses of the key genes in an ER stress response in EA.hy926 cells stimulated by 1 ug/mL tunicamycin for 48 h treated with different concentration of KUS121 (50 and 100 μM) or DMSO as control (n=3 per group). **(B)** Representative western blotting images of ER stress-induced apoptosis marker CHOP and BIP, and VCP in EA.hy926 cells cultured with 1 ug/mL tunicamycin for 48 h treated with different concentration of KUS121 (50 and 100 μM) or DMSO as control, and **(C)** quantification by densitometry (n=3 per group). **(D)** Quantitative real-time PCR analyses of the inflammatory genes downstream of the ER stress pathway. EA.hy926 cells were cultured with 1ug/mL tunicamycin for 48 h with different concentration of KUS121 (50 and 100 μM) or DMSO as control (n=3 per group). **(E)** Representative western blotting images of ER stress-induced phosphorylation of IRE1α, JNK and NF-κB in EA.hy926 cells cultured with tunicamycin for 48 h with different concentration of KUS121 (50 and 100μM) or DMSO as control and **(F)** quantification by densitometry (n=3 per group). The experiments were independently repeated 3 times. All data are presented as mean ± SEM. *p<0.05 **p<0.01, ***p<0.001 by 1-way ANOVA with the Tukey’s multiple comparisons test.

The ER stress pathway is also known to induce inflammation, a pivotal player in atherosclerosis.^31,32^ Thus, we hypothesized that the inhibitory effects of KUS121 on ER stress could concomitantly attenuate inflammation, which has never been investigated before. To test this hypothesis, we first assessed the transcriptional changes of the inflammatory genes in EA.hy926 cells under ER stress. Our results revealed that KUS121 treatment significantly counteracted the upregulation of the Intercellular Adhesion Molecule1 (ICAM1), interleukin (IL) 1β (IL1B), IL6, and IL8, driven by tunicamycin-induced ER stress, providing the evidence of its anti-inflammatory effects (Figure 3D). Since the Inositol-requiring enzyme (IRE) 1α-c-Jun N-terminal kinase (JNK)-Nuclear factor kappa B (NF-κB) signaling is the main inflammatory pathway induced by ER stress.^31^ We assessed the phosphorylation of these proteins as the markers of their activation. Tunicamycin induced IRE1α phosphorylation, which in turn triggered the phosphorylation of downstream JNK and NF-κB, and KUS121 successfully suppressed the activation of this inflammatory pathway under ER stress (Figure 3E and F). Taken together, KUS121 is thought to be able to reduce the ER stress-induced endothelial apoptosis and inflammation.

On the other hand, we also observed some ER stress responses in plaque macrophages (Figure 1H). Thus, we next assessed the effect of KUS121 on macrophages. However, KUS121 did not affect the viability of the THP-1 derived macrophages and murine primary peritoneal macrophages under tunicamycin-induced ER stress (Figure S3A). Moreover, KUS121 could not attenuate the ER stress-induced CHOP expression in THP-1 derived macrophages (Figure S3B).

### KUS121 can reduce ER Stress induced apoptosis and inflammation at endothelium in vivo

To validate the impact of KUS121 on endothelial cells *in vivo*, we first assessed the endothelial CHOP expression in atherosclerotic plaques of the KUS121-treated group by immunostaining and found that KUS121 treatment significantly reduced the CHOP-positive endothelial cells, as compared to the control group (Figure 4A). Since CHOP induces apoptosis, we also assessed apoptotic endothelial cells in plaques by TUNEL staining and confirmed that KUS121 reduced apoptosis in plaque endothelial cells (Figure 4B). Next, to verify the anti-inflammatory effect of KUS121 on endothelial cells, we investigated the impact of KUS121 on the translocation of NF-κB to the nuclei in plaque endothelial cells. Consequently, KUS121 treatment significantly reduced the nuclear translocation of NF-κB in endothelial cells during atherosclerosis progression (Figure 4C). We also demonstrated that ICAM1 which is the critical adhesion molecule for recruitment of monocytes was significantly reduced in plaque endothelial cells of the KUS121-treated group, as compared to the control group (Figure 4D). Consistent with this result, monocyte recruitment into atherosclerotic plaques assessed by counting bead-labeled monocytes in the plaques was also reduced in KUS121-treated group, as compared to the control group (Figure 4E). The reduction of peripheral neutrophils and Ly6C^hi^ inflammatory monocytes may also contribute to the reduction of monocyte recruitment (Figure S1). These data further corroborate that KUS121 can attenuate atherosclerosis progression by reducing ER stress-induced endothelial apoptosis and inflammation.

**Figure 4.**
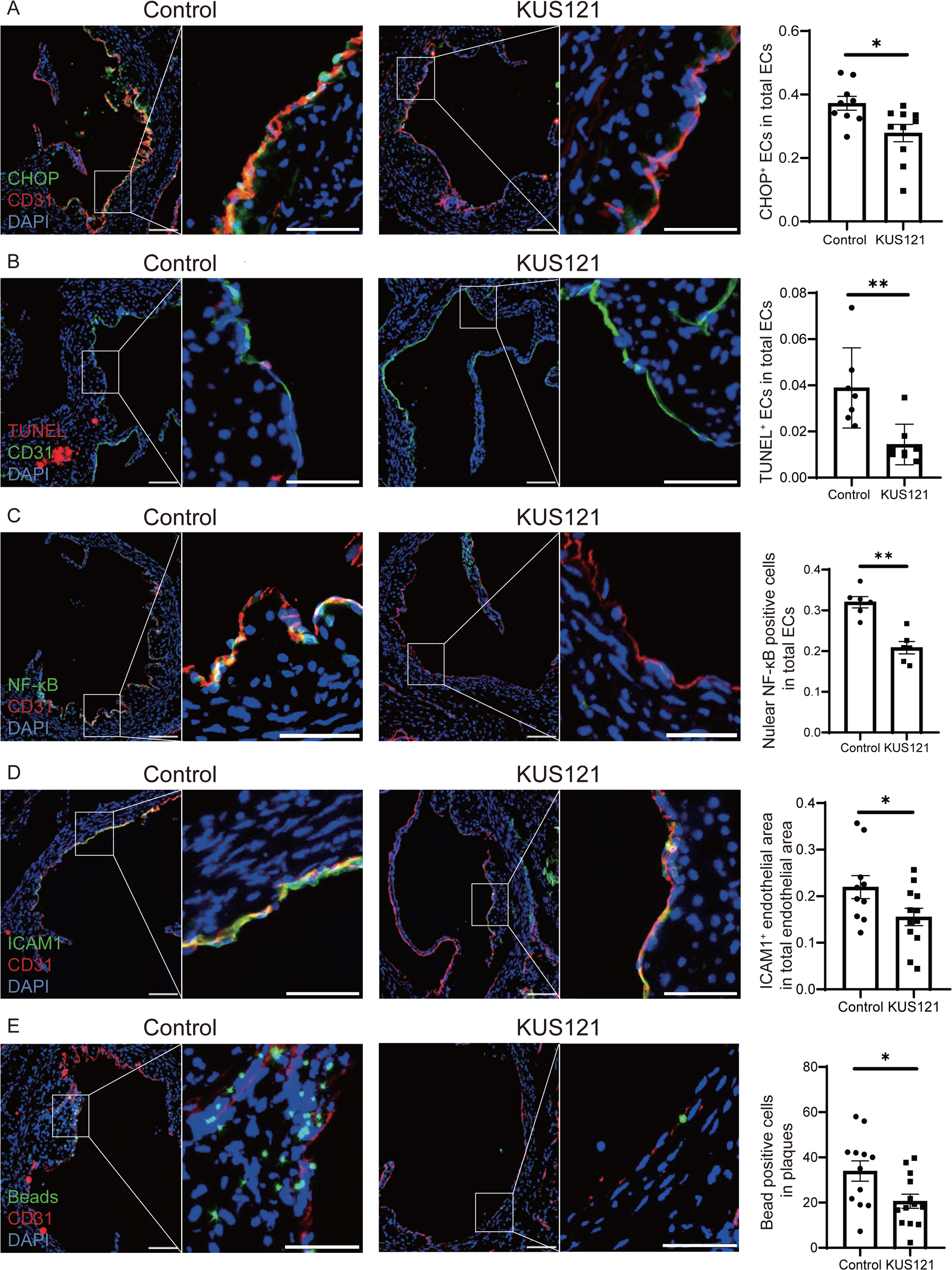
KUS121 reduced ER stress-induced endothelial apoptosis and inflammation *in vivo*. **(A)** Representative endothelial CHOP expression in the aortic sinuses of the control group (left) and the KUS121-treated group (middle), and the proportion of CHOP-positive endothelial cells in the total endothelial cells (right) (n=9-10 per group). *p<0.05 by Mann-Whitney *U* test. **(B)** Representative TUNEL-positive cells in the aortic sinuses of the control group (left) and the KUS121-treated group (middle), and the proportion of TUNEL-positive endothelial cells in the total endothelial cells (right) (n=7-8 per group). **p<0.01 by student’s *t* test. **(C)** Representative endothelial NF-κB expression in the aortic sinuses of the control group (left) and the KUS121-treated group (middle), and the proportion of nuclear NF-κB-positive endothelial cells in the total endothelial cells (n=6 per group). **p<0.01 by student’s *t* test. **(D)** Representative endothelial ICAM1 expression in the aorta sinuses of the control group (left) and the KUS121-treated group (middle), and the proportion of the ICAM1-positive area in the total endothelial area. (n =10-12 per group). *p<0.05 by student’s *t* test. **(E)** Representative bead-labeled cells in the aortic sinuses of the control group (left) and the KUS121-treated group (middle), and the number of bead-labeled cells in atherosclerotic plaques (n=12-13 per group). *p<0.05 by student’s *t* test. All data are presented as mean ± SEM. Scale bar of low magnification images: 200 μm and high magnification images: 50 μm.

### KUS121 can maintain intracellular ATP levels under ER stress and counteract ER stress response in endothelial cells

KUS121 is also known to maintain intracellular ATP levels in several pathological conditions.^13,21,22^ To confirm this effect in endothelial cells, we employed the fluorescent ATP sensor, iATPSnFR^1.0^ by which cytosolic ATP levels can be assessed as a fluorescence intensity of GFP.^28^ As a result, KUS121 successfully rescued the reduction of intracellular ATP under ER stress induced by tunicamycin (Figure 5A-B). KUS121 also reverted the ATP reduction by the treatment of 2DG (Figure S3C-D). Extracellular ATP is reported to induce vascular inflammation to promote atherosclerosis via P2Y2 receptor.^33^ However, it is still unclear how intracellular ATP levels affect atherosclerosis progression especially in endothelial cells. Since an ER stress consumes intracellular ATP (Figure 5A-B) and KUS121 was designed to inhibit ATPase activity of VCP to save intracellular ATP, it is thought that the attenuation of ER stress by KUS121 is due to the maintenance of intracellular ATP levels although it is not fully elucidated. To confirm this, we adopted methyl pyruvate (MePyr) to increase intracellular ATP levels specifically, which is a membrane-permeable pyruvate and known to be converted to ATP in mitochondria. It actually increased intracellular ATP levels in endothelial cells under tunicamycin-induced ER stress (Figure 5C). Moreover, we found that MePyr treatment successfully attenuated ER stress-induced apoptosis and the activation of inflammatory pathways (Figure 5D-E). On the other hand, addition of extracellular ATP did not reduce ER stress responses (Figure S3E), which indicates that intracellular and extracellular ATP work differently for endothelial cells.

**Figure 5.**
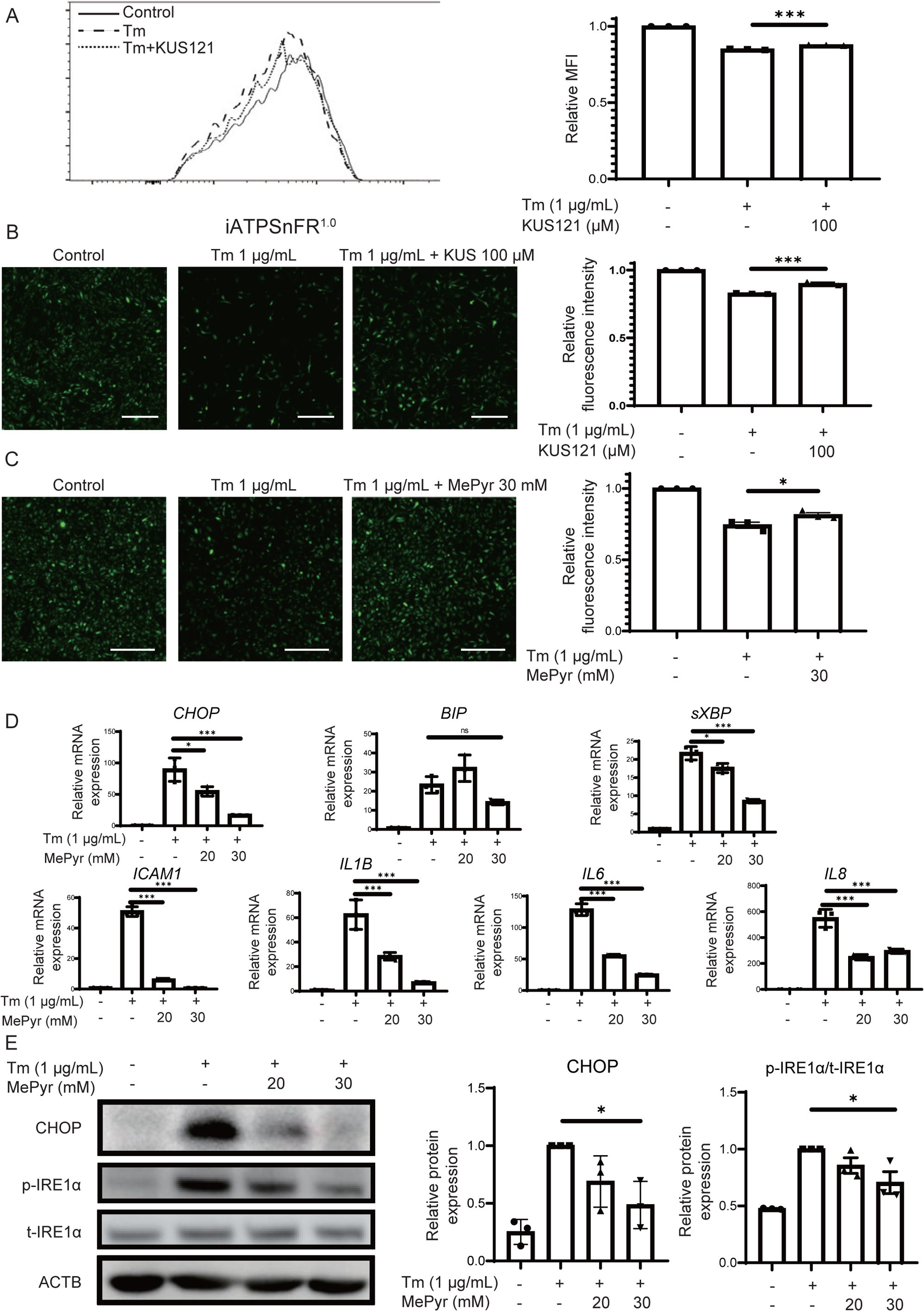
KUS121 maintained endothelial ATP levels under ER stress, which attenuated ER stress responses *in vitro.* **(A)** Representative histogram of the fluorescence intensity of the iATPSnFR^1.0^ vector transfected into EA.hy926 cells cultured with 1 μg/mL tunicamycin for 6 h with or without 100 μM of KUS121 (left) and quantification of relative mean fluorescence intensity (right) (n=3 per group). **(B)** Representative microscopic fluorescence images of the iATPSnFR^1.0^ transfected EA.hy926 cells cultured with 1 μg/mL tunicamycin for 48 h with or without 100 μM KUS121 (left) and quantification of relative fluorescence intensity (right) (n= 3 per group). Scale bar: 50 μm. **(C)** Representative microscopic fluorescence images of the iATPSnFR^1.0^ transfected EA.hy926 cells cultured with 1 μg/mL tunicamycin for 24 h with or without 30 mM Methyl Pyruvate (MePyr) (left) and quantification of relative fluorescence intensity (right) (n= 3 per group). Scale bar: 50 μm. **(D)** Quantitative real-time PCR analyses of the key genes in an ER stress response and the inflammatory genes downstream of the ER stress pathway in EA.hy926 cells cultured with 1 ug/mL tunicamycin for 24 h supplemented with different concentration of MePyr (20 and 30 mM) or DMSO as control (n=3 per group). **(E)** Representative western blotting images of CHOP and phosphorylation of IRE1α in EA.hy926 cells cultured with 1ug/mL tunicamycin for 24 h with different concentration of MePyr (20 and 30 mM) or DMSO as control (left) and quantification by densitometry (right) (n=3 per group). The experiments were independently repeated 3 times. All data are presented as mean ± SEM. *p<0.05, ***p<0.001 by 1-way ANOVA with the Tukey’s multiple comparisons test.

### KUS121 can restore ATP levels of atherosclerotic plaque lesions

Next, we tried to confirm that KUS121 can increase ATP levels at atherosclerotic plaques. To assess ATP levels *in vivo*, we employed GO-ATeam2 *Apoe^−/−^* mice which carry genetically encoded Förster Resonance Energy Transfer (FRET) biosensor.^22^ The FRET ratios at atherosclerotic lesions in the common carotid arteries were calculated pre and post intraperitoneal injection of KUS121 or 5 % glucose as control at several time points. Compared to the control group, KUS121 maintained significantly higher ATP levels at atherosclerotic lesions (Figure 6).

**Figure 6.**
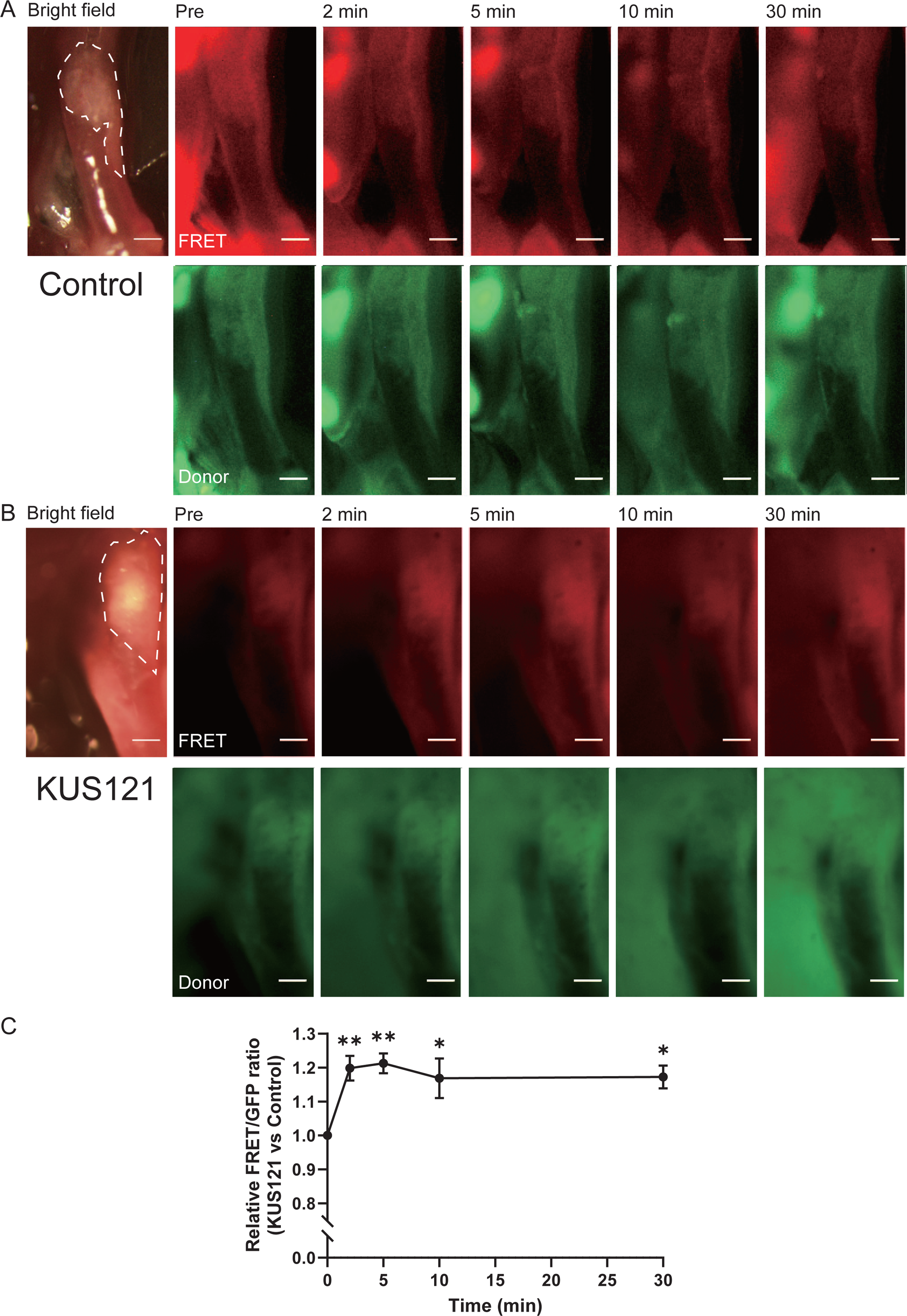
KUS121 increased the ATP levels at atherosclerotic plaque lesions. Representative time course fluorescence images in FRET (upper) and donor (lower) channels at atherosclerotic plaque lesions of common carotid arteries (plaques are indicated by dashed lines) in Go-Ateam2 *Apoe^−/−^* mice on WD for 12 months pre and post **(A)** 5 % glucose solution as control and **(B)** KUS121 (50mg/kg) administration. **(C)** The ratio of FRET/GFP values between KUS121 and control groups at each timepoint (n=3 per group). Scale bar: 200 μm. All data are presented as mean ± SEM. *p<0.05 **p<0.01 by 2-way ANOVA with the Tukey’s multiple comparisons test.

## Discussion

We had previously demonstrated that KUS121 attenuated ER stress-induced apoptosis in several disease models.^13,21,22^ Given the unfavorable effect of ER stress on atherosclerosis,^29^ it was expected that KUS121 can mitigate atherosclerosis. Actually, the daily treatment of KUS121 reduced atherosclerosis progression by approximately 40%. It has been thought that ER stress responses in all major cell populations in atherosclerotic plaques, such as macrophages, endothelial cells and smooth muscle cells contribute to the exacerbation of atherosclerosis progression although less is known about smooth muscle cells.^4,29,34^ Interestingly, CHOP, an established ER stress marker, was mainly expressed in plaque endothelium in our early murine atherosclerosis model. We also observed some CHOP expression in plaque macrophages, but KUS121 could not affect the ER stress in the macrophage at all. Another report showed that ER stress was partly activated in vascular smooth muscle cells during atherosclerosis progression. ^35^ However, we could hardly detect CHOP expression in smooth muscle cells in our atherosclerosis model. Then, we focused on the effect of KUS121 on endothelial cells.

The onset of atherosclerosis involves the injury to the vascular endothelial cells, which can provoke endothelial apoptosis. Numerous conventional risk factors such as oxidized low-density lipoprotein (ox-LDL), low shear stress, etc. are known to induce endothelial apoptosis.^29^ Thus, apoptosis of endothelial cells could be an incipient event in the pathogenesis of atherosclerosis, fostering the initiation and progression of atherosclerotic lesions. Therefore, anti-apoptotic effect of KUS121 was initially considered as a potential reason for the attenuation of atherosclerosis progression by KUS121 treatment. Indeed, we successfully demonstrated that KUS121 can protect endothelial cells from ER stress-induced apoptosis both *in vitro* and *in vivo*.

Atherosclerosis is essentially a chronic inflammatory disease, and inflammation is thought to fuel atherosclerosis progression. This concept was further supported by the recent clinical studies of inhibiting IL-1β by Canakinumab (CANTOS) or administrating colchicine (COLCOT and LoDoCO2). Those anti-inflammatory therapies did not lower atherogenic lipids but reduce cardiovascular events.^36–38^ An ER stress response is also known to activate inflammatory signaling pathways. Specifically, IRE1, one of the ER stress sensors, induces inflammatory responses by interacting TRAF2 to activate NF-κB which is a principal transcriptional regulator of pro-inflammatory pathways. In addition to the activation of NF-κB, the IRE1α-TRAF2 complex can also drive inflammation through the JNK pathway.^31^ Especially, the NF-κB pathway is believed to play a central role in the inflammation of endothelial cells during atherosclerosis progression.^39^ Even though previous studies have demonstrated that KUS121 protect cells from apoptosis by an ER stress response,^13,18,22^ it was still unclear if KUS121 can also attenuate ER stress-induced inflammatory pathways. In this study, we demonstrated that KUS121 can mitigate the ER stress-induced IRE1 pathway, which subsequently reduces the activation of the NF-κB and JNK pathways, and the downstream inflammatory gene expressions. It is notable that the reduced expression of ICAM1, a cell adhesion molecule, in plaque endothelium by KUS121 treatment may account for the reduction of monocyte recruitment into atherosclerotic plaques. We also observed a lower count of peripheral neutrophils and inflammatory Ly6C^hi^ monocytes in the KUS121-treated group, which might be due to the attenuation of systemic inflammation. This reduction of Ly6C^hi^ monocytes in circulation may also devote to the less recruitment of monocytes into plaque.

Original function of KUS121 is to inhibit ATPase activity of VCP to save intracellular ATP consumption.^13,21,22^ We confirmed this effect also in endothelial cells at single cell levels *in vitro* by employing a novel fluorescent ATP sensor, iATPSnFR^1.0^.^28^ It was reported that the depletion of ATP occurred in deep position of plaques coupled with glucose and oxygen deficiency using fresh plaque sections.^40^ In this study, we used a FRET based animal model (Go-Ateam2 mouse) to assess the ATP levels at atherosclerotic lesions *in vivo* and successfully observed KUS121 maintained intracellular ATP levels at the plaque lesions, although it remains uncertain whether the ATP variations in plaque endothelium are consistent with the overall plaque lesion. Administration of extracellular ATP was reported to promote the progression of atherosclerosis via P2Y2 receptor.^33^ However, it still remains to be elucidated how intracellular ATP levels affect atherosclerosis progression especially in endothelial cells. Several steps in an ER stress response require ATP as a source of energy^41^ and we actually observed that intracellular ATP levels were reduced under ER stress at single cell levels, which led to think the idea that KUS121 reduces ER stress by keeping intracellular ATP levels. To clarify this idea, we used methyl pyruvate to selectively increase intracellular ATP levels, and confirmed that maintenance of intracellular ATP levels under ER stress-inducing conditions inhibited both apoptotic and inflammatory pathways of ER stress responses in endothelial cells. These effect of KUS121 could lead to attenuation of atherosclerosis progression. On the other hand, however, addition of ATP extracellularly did not reduce apoptotic or inflammatory gene expressions, rather tended to increase them, which is consistent with the previous reports.^33,42^ Thus, maintenance of intracellular ATP levels in endothelial cells by KUS121 can be the key mechanism for how ER stress is reduced by KUS121 treatment, which in turn mitigates atherosclerosis progression (Figure 7).

**Figure 7.**
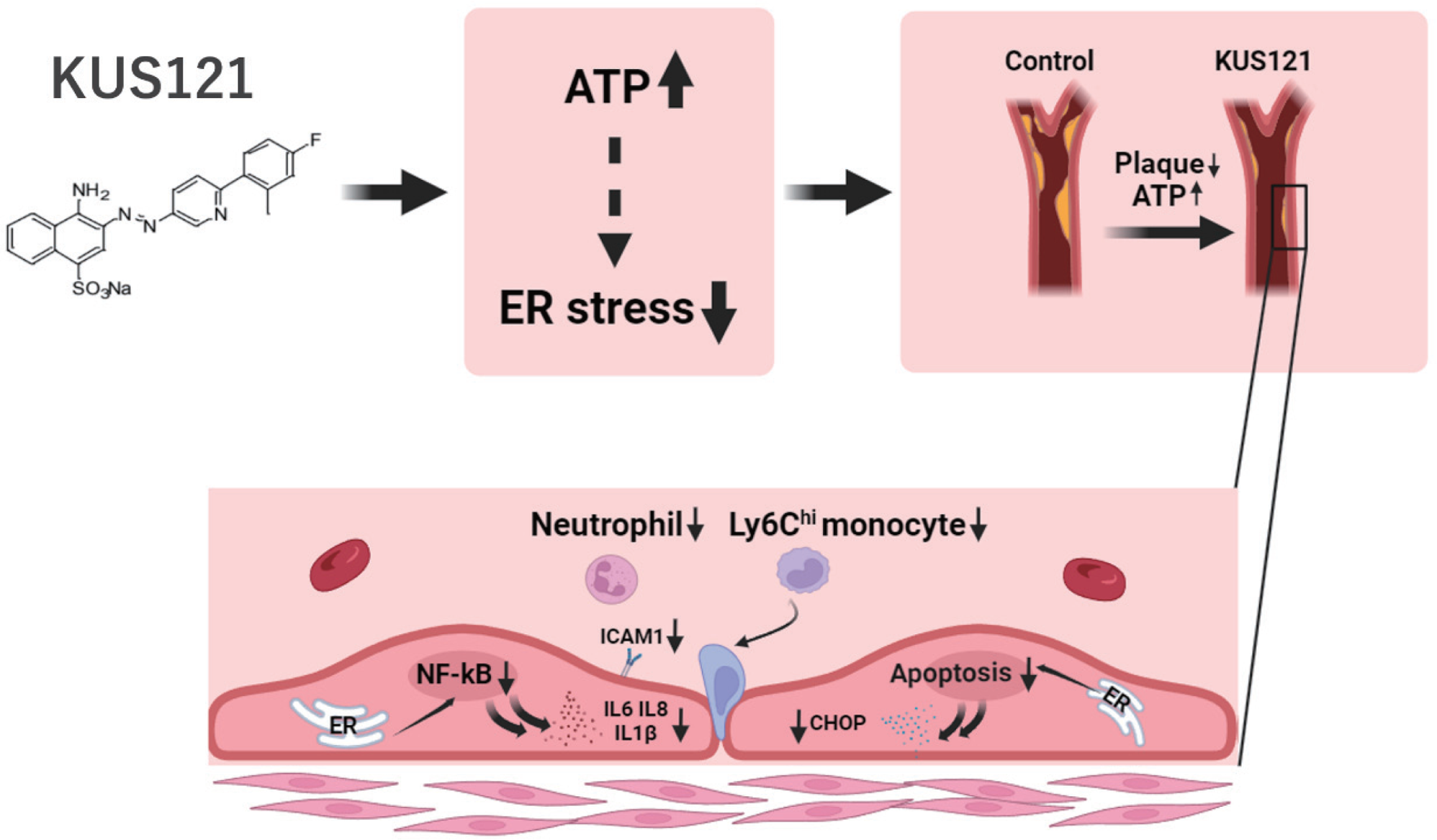
The mechanism how KUS121 attenuates atherosclerosis.

The first limitation of this study is that the off-target effect and cell specificity of KUS121 have not been completely elucidated. In the previous studies, KUS121 consistently protected cells,^13,21,22^ but in the present study, we found that KUS121 did not protect macrophages from ER stress-induced apoptosis. It may be simply because a consumption of ATP by VCP or a response against a maintenance of intracellular ATP levels under ER stress is different in each cell population, but it is also possible that KUS121 might have the off-target effect working just in macrophages. Secondly, the effect of KUS121 on smooth muscle cells might also contribute to a reduction of atherosclerosis, although it may be relatively small based on the less expression of CHOP in smooth muscle cells. Moreover, it’s also possible that ER stress responses occur more in smooth muscle cells at a different time point. Further studies are necessary to answer those questions.

In conclusion, our findings indicate that KUS121 can provide a new therapeutic option for atherosclerosis by reducing ER stress-induced endothelial apoptosis and inflammation, although further studies are needed toward the future clinical application. There are several other ER stress repressors, such as 4-phenyl butyric acid (PBA) and tauroursodeoxycholic acid (TUDCA), and some of them are thought to be used for atherosclerotic disease^4^. However, their mechanisms are different from KUS 121 that specifically inhibits ATPase activity of VCP without affecting other cellular functions. Thus, it is possible that KUS121 can be used with other ER stress repressors to treat atherosclerotic disease, which might be rather synergistically. Moreover, previously we demonstrated that KUS121 has protective effects against ischemic heart disease and heart failure. Those diseases are often comorbid with atherosclerosis. Therefore, KUS121 has a potential to treat those diseases simultaneously.

## Nonstandard Abbreviations and Acronyms

BIP: Binding Immunoglobulin Protein
CHOP: C/EBP homologous protein
ICAM: Intercellular adhesion molecule
IRE: Inositol-requiring enzyme
JNK: c-Jun N-terminal kinase
KUS: Kyoto University substance
MePyr: Methyl pyruvate
NF-κB: Nuclear factor kappa B
VCP: Valosin-containing protein

## Acknowledgements

Imaging analyses including *in vivo* ATP measurements were performed with the support of Kyoto University Live Imaging Center and the Medical Research Support Center, Graduate School of Medicine, Kyoto University.

## Sources of Funding

This work was supported by grants from Japan Society for the Promotion of Science KAKENHI (20K17115 and 22K08154 to O.B., 20K21600 and 23H0294 to K.O.), Uehara Memorial Foundation (to O.B.) and Mochida Memorial Foundation (to O.B.).

## Disclosures

A.K. owns stock in Kyoto Drug Discovery & Development, a start-up company for the development of VCP modulators. This study is partially supported by Kyoto Drug Discovery & Development. The funder has no role in study design, data collection and analysis, decision to publish, or preparation of the manuscript.

## Author contributions

F.Z. and O.B. conceived the main conceptual ideas and designed the research plan. F.Z. conducted the experiments and collected the data. N.S. provided the technical assistance for the experiments. F.Z. and O.B. analyzed and interpreted the data. T.H., Y.N., S.T., T.Y., C.O., S.X., M.I., K.M., K.S., E.K., H.K., Q.Q., K.K. and K.O. aided in interpreting the results. A.K. and K.O. supervised the project. O.B. provided direction throughout the preparation of this manuscript. F.Z. and O.B. wrote the manuscript with support from T.H., A.K. and K.O. All authors read and approved the final manuscript.

## Notes

### Competing Interest Statement

An author owns stock in Kyoto Drug Discovery & Development, a start- up company for the development of VCP modulators. This study is partially supported by Kyoto Drug Discovery & Development. The funder has no role in study design, data collection and analysis, decision to publish, or preparation of the manuscript.

